# The tea plant (*Camellia sinensis*) *FLOWERING LOCUS C-like* gene *CsFLC1* is correlated with bud dormancy and triggers early flowering in *Arabidopsis*

**DOI:** 10.1101/2022.09.02.506372

**Authors:** Ying Liu, Ludovico Dreni, Haojie Zhang, Xinzhong Zhang, Nana Li, Kexin Zhang, Taimei Di, Lu Wang, Yajun Yang, Xinyuan Hao, Xinchao Wang

**Author notes:** Correspondence: Xinchao Wang, Xinyuan Hao, Authors to whom correspondence should be addressed.

## Abstract

Flowering and bud dormancy are crucial stages in the life cycle of perennial angiosperms in temperate climates. MADS-box family genes are involved in many plant growth and development processes. Here, we identified 3 *MADS-box* genes in tea plant belonging to the *FLOWERING LOCUS C* (*CsFLC*) family. We monitored *CsFLC1* transcription throughout the year and found that *CsFLC1* was expressed at a higher level during the winter bud dormancy and flowering phases. To clarify the function of *CsFLC1*, we developed transgenic *Arabidopsis thaliana* plants heterologously expressing *35S::CsFLC1*. These lines bolted and bloomed earlier than the WT (Col-0), and the seed germination rate was inversely proportional to the increased *CsFLC1* expression level. RNA-seq of *35S::CsFLC1* transgenic *Arabidopsis* showed that many genes responding to ageing, flower development and leaf senescence were affected, and phytohormone-related pathways were especially enriched. According to the results of hormone content detection and RNA transcript level analysis, *CsFLC1* controls flowering time possibly by regulating *SOC1, AGL42, SEP3* and *AP3* and hormone signalling, accumulation and metabolism. Our results suggest that *CsFLC1* might play dual roles in flowering and winter bud dormancy and provide new insight into the molecular mechanisms of *FLC* in tea plants as well as other plant species.

**Highlight:** Three *FLOWERING LOCUS C-like* genes were identified in tea plants, among them *CsFLC1* played dual roles in flowering and winter bud dormancy.

## 1. Introduction

Flowering is an important trait that helps plants transition from the vegetative phase to the reproductive phase and involves many complex changes, including those involving physiological, metabolic and molecular processes (Balanzà *et al*., 2018). Moreover, flowering time is regulated not only by a plant’s intracellular signature but also by environmental factors (Dong *et al*., 2021). Daylength and temperature are two main environmental factors that affect plant reproduction. According to the daylength and temperature, plants can sense the season and whether the time is proper to produce flowers and seeds. The vernalization pathway was proposed to explain how temperate angiosperms avoid blooming during winter. FLOWERING LOCUS C (FLC) is considered a crucial regulator in the vernalization pathway (Sheldon *et al*., 2000). The function of *FLC* differs between ecotypes of *Arabidopsis thaliana*: it represses flowering in late-flowering ecotypes (Michaels and Amasino, 1999), while in early-flowering ecotypes, *FLC* overexpression further delays flowering (Ratcliffe, 2001). In Chinese cabbage (*Brassica rapa* ssp. *pekinensis* (Lour.) Hanelt), there are 3 *FLC* homologues that have lower expression levels in early-flowering varieties (Kim *et al*., 2007). Additionally, in *Eustoma grandiflorum* (Raf.) Shinners, *EgFLC* represses flowering (Nakano *et al*., 2011). FLC was reported to directly bind to the promoter of *SUPPRESSOR OF OVEREXPRESSION OF CONSTANS 1* (*SOC1*) and the first intron of *FLOWERING LOCUS T* (*FT*) to repress expression and delay flowering time (Searle, 2006; Seo *et al*., 2009). *FLC* expression is reduced by DNA methylation (epigenetic modifications) but induced by acetylation of histones (chromatin remodelling) (Sheldon *et al*., 1999; He and Amasino, 2005; Greb *et al*., 2007; Sheldon *et al*., 2008; Xiao J *et al*., 2013; Kwak *et al*., 2016; Nishio *et al*., 2016; Questa *et al*., 2016; Kwak *et al*., 2017).

Bud dormancy is necessary for temperate perennials to avoid cellular damage and ensure plant survival during winter. *FLC* was also related to bud dormancy of perennial plant species such as apple (*Malus × domestica* Borkh.) and kiwifruit (*Actinidia chinensis* Planch.), where it is highly expressed during dormancy (Soichiro *et al*., 2019; Voogd *et al*., 2022). However, transgenic kiwifruit plants overexpressing *AcFLCL* displayed earlier budbreak times (Voogd *et al*., 2022). Tea plant (*Camellia sinensis* (L.) Kuntze) is an economically important crop species in many countries and areas of Asia, Africa and Latin America (Wang *et al*., 2020). Its leaves are processed into different kinds of tea, which is famous for its health benefits (Yi *et al*., 2019; Williams *et al*., 2019; Malar *et al*., 2020; Xu *et al*., 2020). Since tea plant is a crop species grown for its leaves, its reproductive phase from flowering to fruit production requires an abundance of plant resources, thus affecting tea plant bud growth and limiting the production and quality of tea. Therefore, breeding improved tea varieties or finding a way to balance vegetative and reproductive growth to improve the efficiency of the tea industry is urgently needed. For this, it is necessary to understand the molecular mechanisms of flowering in tea plant. Tea plant *FLC* orthologues have not yet been described. In our previous study, the phenotypes of two types of tea plant cultivars, a floriferous type and an oliganthous type, were observed (Liu *et al*., 2020). We found that the meristem was maintained in the vegetative phase and did not switch to reproductive growth in the axillary buds of the oliganthous cultivars, while the transition to the floral meristem occurred in June in the axillary buds of the floriferous tea plant cultivars. The flowering-related genes in these two cultivars were identified, and among them, one MADS-box gene was found that might play an important role in floral organ differentiation and maturation (Liu *et al*., 2020). In temperate areas, tea plant usually undergoes bud dormancy in the winter to overcome low-temperature stress, and when the temperature arises in the spring, the buds emerge. The progression of the bud dormancy–budbreak phase involves many genes (Wang *et al*., 2014; Hao *et al*., 2017; Chen *et al*., 2021). Despite several studies about bud dormancy in tea plant, the role of *CsFLC* in dormancy is still unknown. Hence, it is necessary for breeders to understand the mechanisms of flowering and dormancy as well as their relationship.

In this study, we identified 3 *FLC-like* genes in tea plant, among them *CsFLC1*, whose expression was correlated with bud dormancy and flower bud development. The RNA transcription of *CsFLC1* was monitored throughout the year. Then, we established *CsFLC1*-overexpressing transgenic *Arabidopsis* lines to characterize its functionality in a heterologous model plant species. Our study provides a new understanding of flowering as well as bud dormancy in tea plant and a theoretical basis for future breeding programmers to develop novel cultivars with early bud break and few flowers.

## 2. Materials and methods

### 2.1 Plant materials and growth conditions

Buds, leaves, flowers, stems and roots of Longjing 43 (LJ43) and buds of Zhenghedabai (ZHDB) tea plant cultivars for gene expression or hormone detection were collected from a tea plantation at the Tea Research Institute of the Chinese Academy of Agricultural Sciences (Hangzhou, China; N30°180’, E120°100’). Axillary buds located at the same positions along the branches from more than 30 individual plants were collected in the afternoon on Sep. 30^th^, Oct. 14^th^, Nov. 1^st^, and Dec. 16^th^, 2016, and on Jan. 16^th^, Feb. 10^th^, Mar. 1^st^, Mar. 14^th^, Mar. 27^th^, Apr. 13^th^, Apr. 28^th^, May 16^th^, May 27^th^, Jun. 15^th^, Jun. 28^th^ and Jul. 18^th^, 2017. Flower buds were collected in the afternoon on Aug. 15^th^, Aug. 28^th^, Sep. 18^th^ and Sep. 26^th^, 2017. The apical, axillary and floral buds as well as the leaves, flowers, stems and roots of LJ43 were collected on Apr. 12^th^, 2019, for tissue expression analysis. There were three biological replicates collected at each time point. All *Arabidopsis thaliana* plants except those composing the treatment groups were grown under long days (LDs; 16 h light/8 h darkness) at 22 °C and under 70% relative humidity.

Tissues of *Arabidopsis thaliana* for RNA extraction were frozen in liquid nitrogen immediately after harvesting and stored at -80 °C before use.

### 2.2 Phylogenetic analysis

The two tea plant genome assemblies, namely, GCA_004153795.1 (Wei *et al*., 2018) and GCA_013676235.1 (Zhang *et al*., 2020), available in the NCBI database were screened to identify the *FLC* genes and their collinear genes. Only the latter assembly was used to study conserved gene collinearity because of its high continuity at the chromosome-scale level. The other genes and genomes were accessed through Phytozome v.13; several incomplete annotations were found that we corrected by screening the NCBI GenBank database. Gene collinearity was assessed by SynFind (https://genomevolution.org/coge/SynFind.pl) and manually.

To construct a phylogenetic tree, SBP proteins were aligned using MAFFT (https://mafft.cbrc.jp/alignment/server/) and then analysed with MEGA 11 (Tamura *et al*., 2021). The evolutionary history was inferred by using the maximum likelihood method and the Jones Taylor Thornton (JTT) matrix-based model. The trees were drawn to scale, with branch lengths equal to the number of substitutions per site.

The accessions used are listed in Figure 1 and in Supplementary Table S1.

**Fig. 1.**
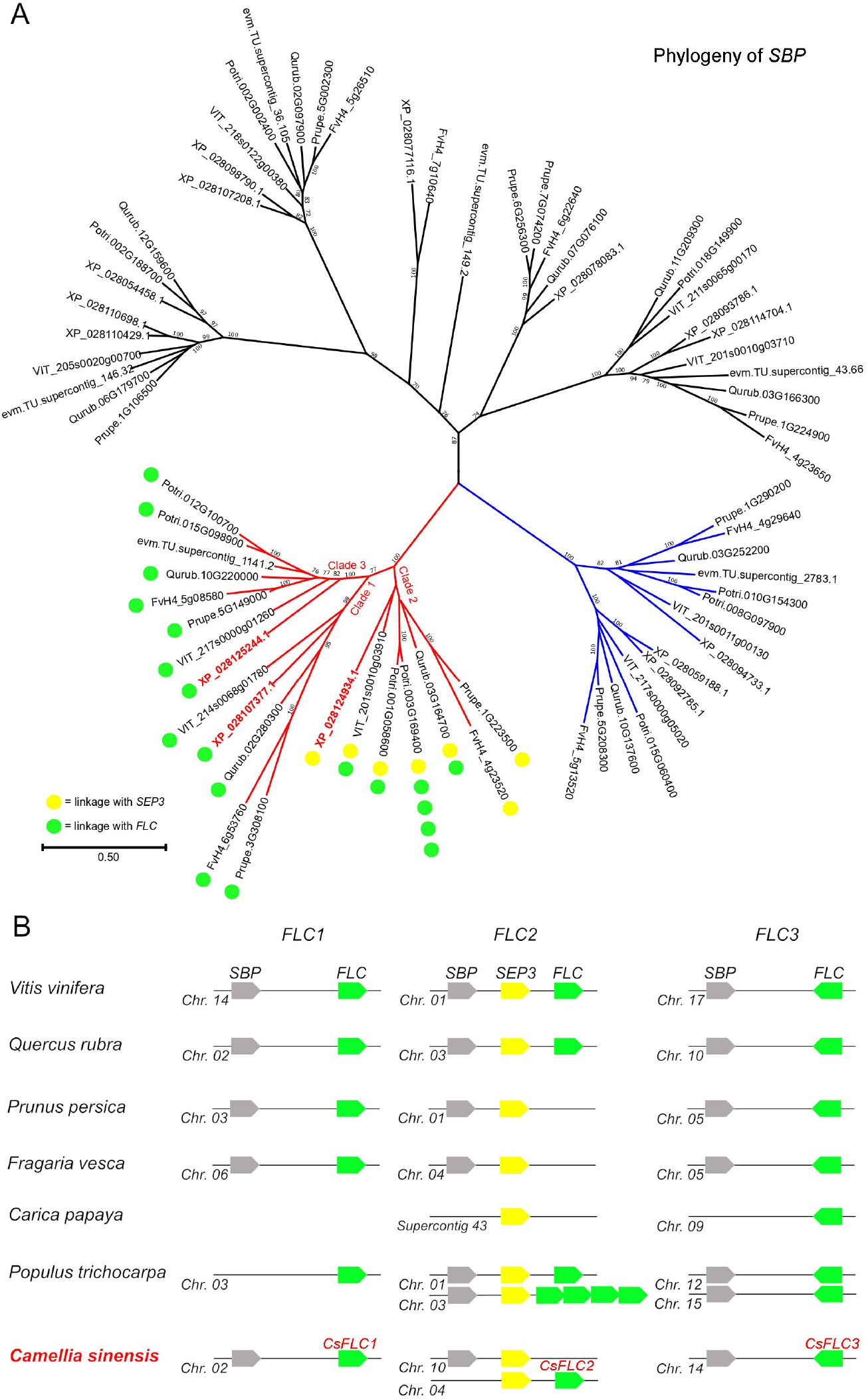
Identification and characterization of the *CsFLC1* gene. A. As reported earlier by Rudall *et al*. (2013), *SBP* genes from a conserved monophyletic clade, shown in red, are closely linked to *SEP3* and *FLC*, and they are triplicated in core eudicots (clades 1, 2 and 3). Two of the three *SBP* genes that we found in *Camellia sinensis* show this conserved linkage with *FLC* genes. The genes of sister clade shown in blue are instead linked to *LOFSEP* and *SQUA* MADS-box genes. Potri: *Populus trichocarpa*; VIT: *Vitis vinifera*; XP: *Camellia sinensis*; Prupe: *Prunus persica*; FvH4: *Fragaria vesca*; Qurub: *Quercus rubra*; evm: *Carica papaya*. The tree was generated using protein sequences. Bootstrap values lower than 70 are not shown, and the scale bar indicates the number of amino acid sequence substitutions per site. (B). The three *SBP*-(*SEP3*)-*FLC* paralogous tandems of core eudicots show different patterns of conservation and duplications among species. In *Camellia sinensis*, which has an ancient tetraploid genome (Chen *et al*., 2020), six regions are expected; however, only duplicated copies of *SEP3* were retained. The results of this analysis suggest the presence of only three *FLC* genes in tea plant, which we refer to as *CsFLC1, CsFLC2* and *CsFLC3*.

### 2.3 Quantitative real-time PCR (qRT□PCR) analysis

Total RNA was isolated from tissues of tea plant as well as *Arabidopsis thaliana* using an RNAprep Pure Plant Kit (Tiangen Biotech Co., Ltd, China) according to the manufacturer’s protocols. For qRT□PCR, 1 μg of total RNA was used to synthesize first-strand cDNA with PrimeScript RT enzyme together with gDNA eraser (Takara, Japan). qRT□PCR was performed on a Roche LightCycler 480 (Roche Diagnostics, Germany) using LightCycler 480 SYBR Green I Master Reagent (Roche Diagnostics, Germany). The gene-specific primer pairs used are listed in Table S2. The polypyrimidine tract-binding protein (*CsPTB1*) gene was used as an internal control (Hao *et al*., 2014). The relative expression levels were calculated using the 2^−^ΔΔCt method (Schmittgen and Livak, 2008).

### 2.4 Cloning, plasmid construction and transformation

Due to the current lack of efficient tea plant transformation techniques, we used the model plant species *Arabidopsis thaliana* as a heterologous expression system to validate *CsFLC1’s* function. The primers that we designed to clone the promoter and coding DNA sequence (CDS) of *CsFLC1* are listed in Table S2. The CDS of *CsFLC1* was cloned into pCAMBIA1300 (p1300) together with a 35S promoter vector to obtain the *35S::CsFLC1* p1300 plasmid. The 1066 bp region upstream of the ATG codon of *CsFLC1* was cloned into p1300 together with a β-glucuronidase (*GUS*) fragment vector, yielding *pCsFLC1::GUS* p1300 plasmids. We transformed the plasmids into *Agrobacterium tumefaciens* GV3101. Transgenic *35S::CsFLC1* and *pCsFLC1::GUS Arabidopsis thaliana* plants were obtained by the floral dip technique (Bent, 1999). The resulting T_0_ generation seeds were germinated on 15 mg/L hygromycin B selective 1/2-strength Murashige and Skoog (MS) plates to select positive plants from among those composing the T_1_ generation, and then the positive plants were transplanted into soil to collect T_1_ seeds. The T_1_ seeds were sown on selective MS plates as described above to select T_2_-generation seedlings showing a positive:negative selection ratio ≈ 3:1; these seedings were transplanted into soil to obtain T_2_-generation seeds. The T_2_ seeds were plated on selective MS plates as described above to select the lines that were 100% hygromycin-resistant, which constituted T_3_-generation homozygous lines.

### 2.5 Subcellular localization

To obtain *35S::CsFLC1:eGFP* recombination plasmids, we cloned the CDS of *CsFLC1* into p1300 together with the 35S promoter and a *GFP* reporter vector. Then, we transformed the resulting plasmids into *Agrobacterium tumefaciens* GV3101. Transformation of *Nicotiana benthamiana* was performed according to a previous description (Kapila *et al*., 1997). An Olympus FV1000 confocal laser-scanning microscope (Zeiss, Germany) was used for imaging.

### 2.6 GUS staining

For cold treatment, 7-day-old *pCsFLC1::GUS Arabidopsis* transgenic lines were exposed to 4 °C for 0, 3 or 4 h. For photoperiod treatment, *pCsFLC1::GUS* transgenic seedlings were grown under 3 different daylengths: LDs (16 h light/8 h darkness), medium days (MDs; 12 h light/12 h darkness) and short days (SDs; 8 h light/16 h darkness) after germinating for one week. Seedlings were collected at the end of the light or dark phase. Different samples or tissues of transgenic *Arabidopsis thaliana pCsFLC1::GUS* plants were subjected to GUS staining buffer as previously described (Wang *et al*., 2018). Two independent T_3_ homozygous lines were used for GUS staining in each experiment. An Olympus SZ61 microscope and a Nikon Eclipse 80i microscope were used for imaging.

### 2.7 RNA sequencing and transcriptome analysis of Arabidopsis transgenic lines

Leaves from twenty-two-day-old Arabidopsis Col-0 and three different *CsFLC1* overexpression T3 homozygous lines (OE 3-10, OE 5-1 and OE 6-1) were collected to perform RNA sequencing. Twelve plants were pooled as one biological replicate, and three replicates of each line and of ecotype Columbia (Col-0) were used for RNA sequencing. A Venn diagram was generated by TBtools (Chen *et al*., 2020). Gene expression pattern clusters were evaluated via R version 3.6.1. Gene Ontology (GO) enrichment was analysed by GOEAST tools (Zheng and Wang, 2008). Kyoto Encyclopedia of Genes and Genomes (KEGG) functional annotation clustering was performed by using DAVID (Dennis *et al*., 2003).

### 2.8 Measurements of phytohormone contents in tea plant buds

For hormone extraction and content determination, 0.1 g of buds of tea plant that were the same samples as those used for RNA extraction were ground into powder in liquid nitrogen; there were three biological replicates for each time point. The methods that we used to extract and determine the contents of the phytohormones have been described previously (Lu *et al*., 2020). Diagrams of the results were generated by GraphPad Prism 6 software (USA).

## 3. Results

### 3.1 Identification of *CsFLC-like* genes in the tea genome

In our previous work, we isolated a MADS-box gene candidate that might play an important role in floral organ differentiation and maturation (Liu *et al*., 2020). Then, we further analysed its putative protein product and found that it is most homologous to *FUL*-like and *FLC*-like genes. Considering its high degree of sequence divergence and to understand its true identity, we applied a mixed approach of phylogeny and gene collinearity. A phylogenomic study (Rudall *et al*., 2013) clarified that one paleo-*FLC* gene was present in the ancestor of angiosperms and that, in fact, *FLC*-like genes also exist in monocots but were previously misclassified. Their conserved genomic locations and their tendency to duplicate only by whole-genome duplication can help overcome the limits of phylogenetic analysis. In particular, the ancestral genomic configuration of *FLC* is in a narrow tandem array together with a monophyletic group of *SQUAMOSA BINDING PROTEIN* (*SBP*) members and with *SEPALLATA 3* (*SEP3*) (Rudall *et al*., 2013). Since core eudicots have paleohexaploid ancestor, this *SBP-SEP3-FLC* cluster should have triplicated in those members. Indeed, in addition to another cluster that we have found on chromosome 17 (Fig. 1A, B) in the model genome of grape (*Vitis vinifera* L.), two such clusters have been reported by Rudall *et al*. (2013). However, *FLC* and its close homologous *MAF* genes lost this conserved collinearity in the *Arabidopsis* family, which is a member of the *Brassicaceae*, probably by gene transposition or a massive genome fractionation (Zhao *et al*., 2017).

We found that the conservation of the three *SBP* paralogous clades is remarkable among core eudicot species (excluding members of the *Brassicaceae*), and most of them reveal an *FLC* locus in close proximity (Fig. 1A, B). In tea plant, two of these *SBP* genes are located near a putative *FLC* locus, and a *SEP3-FLC* tandem array also is present (Fig. 1A, B). Although the tea plant genome underwent a relatively ancient whole-genome duplication event (Chen *et al*., 2020), only duplicated copies of *SEP3* seem to have been retained within the *SBP-SEP3-FLC* clusters, evidence of multiple gene losses (Fig. 1B, Table S1). Finally, a fourth highly diverged MADS-box gene without any clear homology to other core eudicot genes we have screened thus far and in a region noncollinear to *FLC* was isolated by BLAST (GenBank XP_028119527.1), for which further analysis is needed. In conclusion, three paralogous *FLC* clades seem to exist in core eudicots, with one member each in the tea plant genome; we named these genes *CsFLC1, CsFLC2* and *CsFLC3*. Although *CsFLC1* is only 32% homologous to Arabidopsis *FLC*, it is 56% homologous to and collinear with the *AcFLCL* (Acc05562) gene recently reported in the closely related kiwifruit (Voogd *et al*., 2022). Moreover, *CsFLC2* is homologous and shares gene collinearity with the kiwifruit genes Acc14299 and Acc33776 (data not shown).

Since *FLC-like* genes are expressed higher during dormancy in apple and kiwifruit, we analysed the expression patterns of *CsFLC1, CsFLC2* and *CsFLC3* during different dormancy states, including endodormancy, ecodormancy, paradormancy and bud flush, in tea plant via transcriptome analysis (Hao *et al*., 2017),(Fig. S1). The results showed that only *CsFLC1* had a high mRNA expression level during winter dormancy (endo- and ecodormancy), while the expression of *CsFLC2* did not change across the four states, and *CsFLC3* was not expressed at all. Hence, *CsFLC1* was chosen for further analysis.

### 3.2 *CsFLC1* gene expression patterns

To identify the subcellular location of *CsFLC1*, we constructed a *35S::CsFLC1:eGFP* plasmid and transformed it into *Agrobacterium* to infect nuclear marker (red fluorescence) transgenic tobacco. Through confocal microscopy, we observed green fluorescence merged with red fluorescence to show yellow light. As expected for a putative transcription factor (TF), *CsFLC1* was located in the cell nucleus (Fig. 2).

**Fig. 2.**
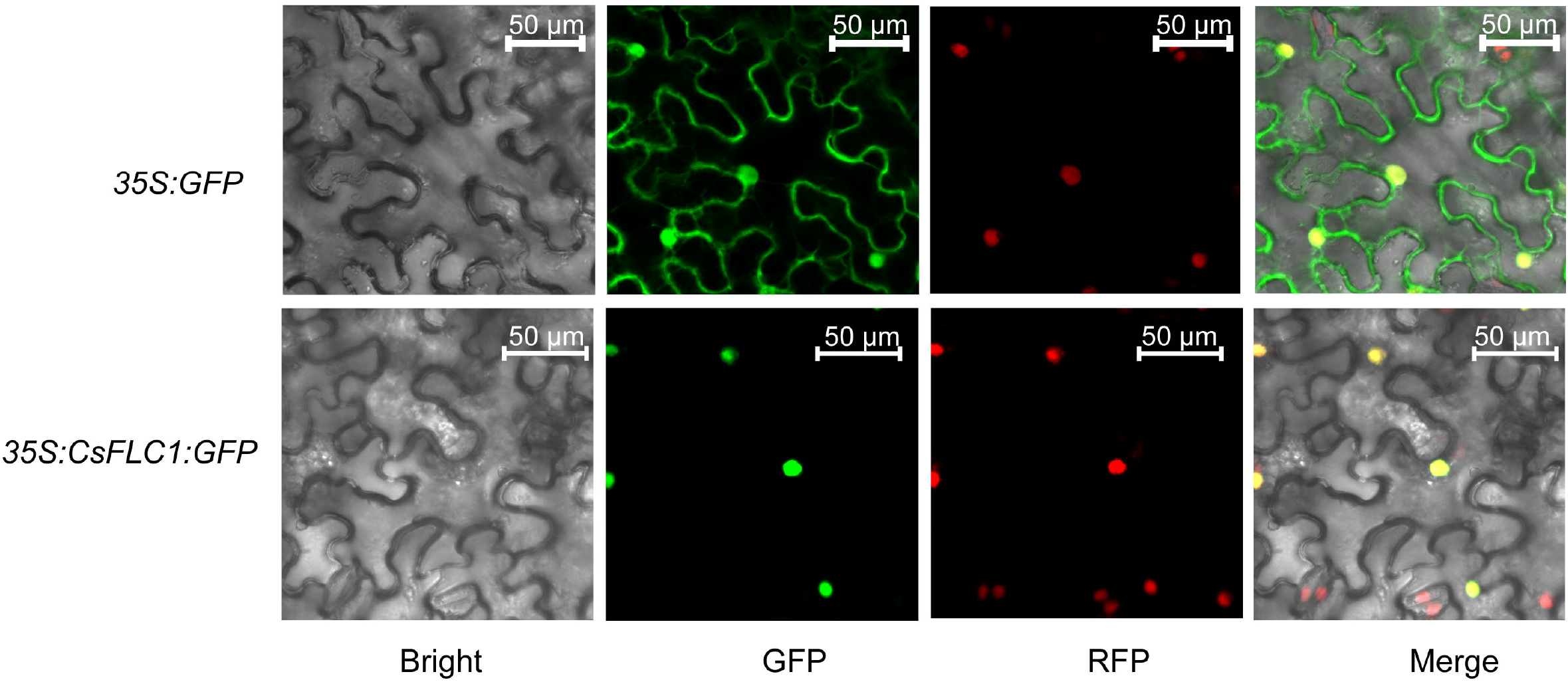
Subcellular location of CsFLC1 in *Nicotiana benthamiana*. GFP: green fluorescent protein; RFP: red fluorescent protein; bar: 50 µm.

To study the expression pattern of *CsFLC1*, we measured the gene expression level in axillary or floral buds in tea plant (Fig. 3A). The results showed that *CsFLC1* had two expression peaks (one on Dec. 28^th^ and another on Aug. 28^th^) throughout the year (from 2016 to 2017). Nov. 1^st^ to the following Mar. 14^th^ corresponded to a dormancy period in tea plant (Hao *et al*., 2017). From May 27^th^ to Sep. 26^th^, floral bud differentiation and floral bud development phases occurred (Liu *et al*., 2020). These results showed that *CsFLC1* expression corresponded to bud dormancy and flowering.

**Fig. 3.**
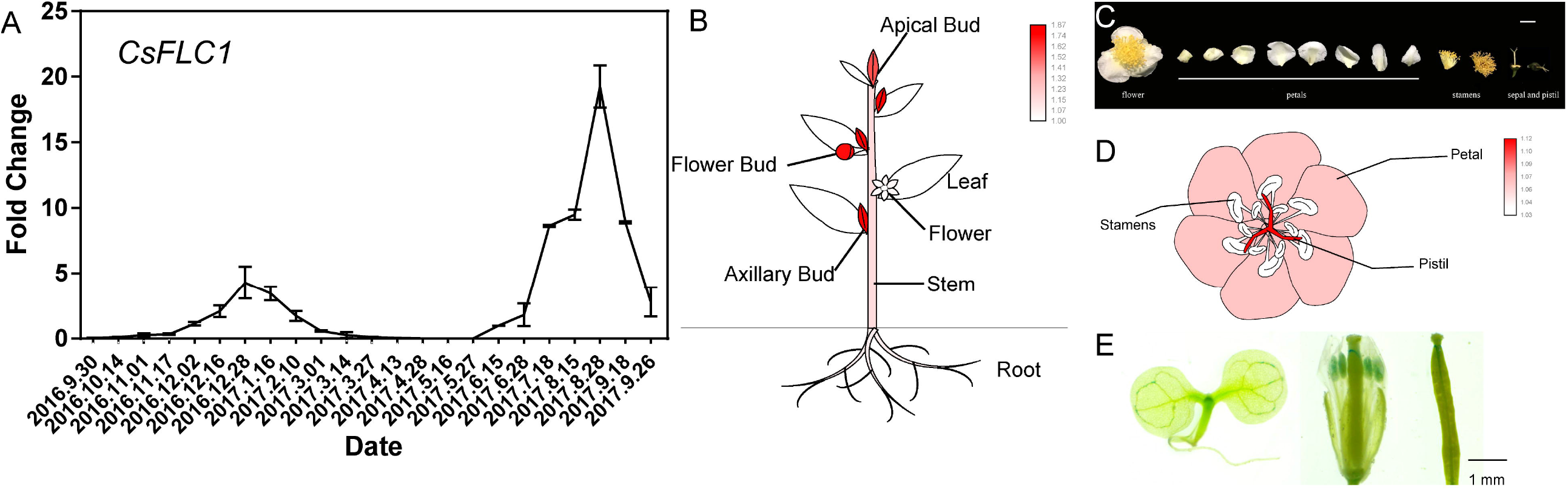
Expression patterns of *CsFLC1*. (A) Expression of *CsFLC1* in axillary or flower buds of tea plant throughout the year. (B) Different tissue expression of *CsFLC* in tea plant. (C) Various parts of flowers of tea plant. (D) Expression patterns of *CsFLC1* in different parts of flowers in tea plant. (E) GUS staining of *pCsFLC1::GUS::GUS* transgenic *Arabidopsis thaliana*.

To identify in which tissues *CsFLC1* is expressed, we measured its expression in seven different tissues of tea plant, namely, apical buds, axillary buds, flower buds, flowers, mature leaves, stems and roots (Fig. 3B). *CsFLC1* had the highest expression level in the three kinds of buds, and the next highest expression was in the stems and roots, while the lowest was in the flowers and leaves. Therefore, we further measured the expression in different flower organs (Fig. 3C). The results showed that *CsFLC1* had the highest expression level in the pistils followed by the petals, while it was not detected in stamens. In *pCsFLC1::GUS::GUS* transgenic *Arabidopsis*, we found similar results (Fig. 3D). *GUS* was expressed in the apical meristems, pistils, and stamens followed by the vascular tissue but was hardly detected in the roots and leaves. The expression patterns in tea and *pCsFLC1:::GUS::GUS Arabidopsis* were common in the apical meristem and pistil but different in the petals, stamens and roots, which might be caused either by an incomplete *pCsFLC1* promoter region or by differences in the *Arabidopsis* heterologous system.

### 3.3 The *CsFLC1* promoter is responsive to low temperature and photoperiod

To clarify the *CsFLC1* response to environmental stimuli, we subjected *pCsFLC1::GUS* transgenic lines to low temperature (4 °C) and different daylengths, namely, LDs (16 h light/8 h darkness), MDs (12 h light/12 h darkness) and SDs (8 h light/16 h darkness), and observed the relative GUS staining patterns. The results showed that the GUS staining in the leaf veins, leaf apexes and roots was stronger under low-temperature treatment than under the control treatment (Fig. 4A, B, C). GUS was expressed at high levels either in the light or dark under the MD treatment (Fig. 4E, H), while it was expressed at higher levels in the light than in the dark under the LD treatment (Fig. 4D, G), and its behaviour was opposite under the SD treatment (Fig. 4F, I).

**Fig. 4.**
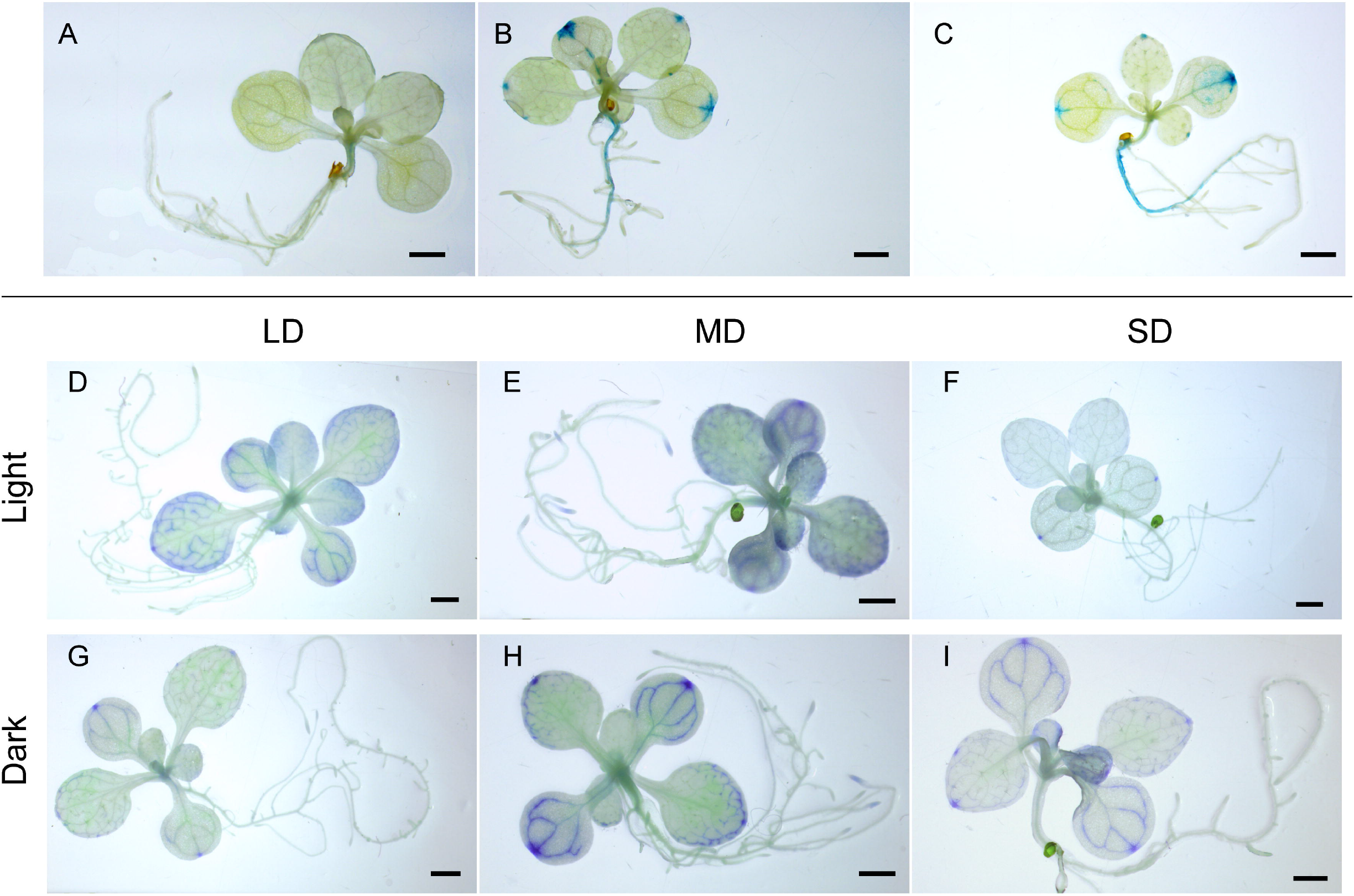
*CsFLC1* in response to low temperature and daylength. (A-C) *pCsFLC1::GUS::GUS* response to low temperature (LT; 4 °C); (A) before LT treatment; (B) after LT treatment for 3 hours; (C) and after LT treatment for 4 h. (D-F) *pCsFLC1::GUS* staining in the light; (D) in plants under LDs; (E) in plants under MDs; and (F) in plants under SDs. (G-I) *pCsFLC1::GUS* staining in the dark; (G) in plants under LDs; (H) in plants under MDs; (I) and in plants under SDs. LD: long days; MD: medium days; SD: short days

### 3.4 *CsFLC1* affects flowering time and seed germination of transgenic *Arabidopsis thaliana*

To determine the function of *CsFLC1*, we tested three independent transgenic of *35S*::*CsFLC1 Arabidopsis* lines (Fig. 5B). We observed the phenotype of the overexpression (OE) lines and found that the bolting, flowering and ageing time of the transgenic lines occurred earlier than did those of Col-0 wild type (WT; Fig. 5A, D). In addition, the higher *CsFLC1* expression was, the lower the seed germination rate was (Fig. 5C).

**Fig. 5.**
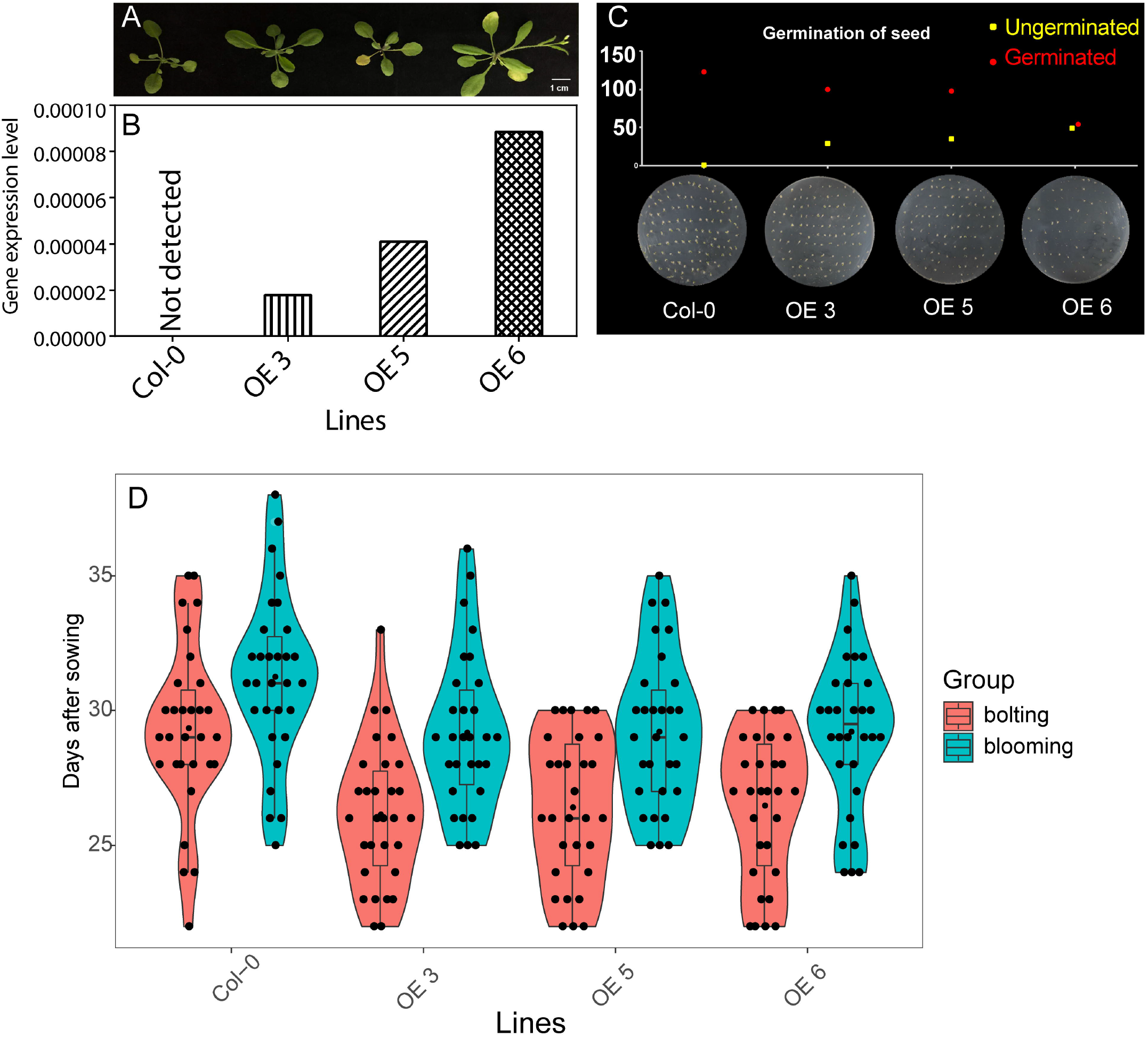
Phenotypes of *35S::CsFLC1* transgenic *Arabidopsis thaliana*. (A) Image of WT and *OE-CsFLC1* lines. (B) *CsFLC1* expression levels in different *Arabidopsis thaliana* lines. (C) Seed germination rates of different *Arabidopsis thaliana* lines. (D) Boxplot showing bolting and blooming times of different *Arabidopsis thaliana* lines.

To further study how *CsFLC1* plays roles in these physiological processes, we measured the gene expression levels of three OE lines and WT *Arabidopsis* via RNA sequencing. In total, 228 differentially expressed genes (DEGs) were common in the three OE lines compared with the WT (Fig. S2). In total, 169 DEGs were upregulated in the OE lines compared with the WT, while 59 DEGs were downregulated (Fig. 6A, B). Among the upregulated gene clusters of OE-*CsFLC1* lines, the terms ‘ageing’, ‘flower development’ and ‘leaf senescence’ among biological process (BP) were enriched. Additionally, the OE-*CsFLC1* lines were senescent, and bolting and blooming occurred earlier than they did in the WT. Therefore, the phenotype and regulated pathways were matched. The genes associated with the terms ‘response to abscisic acid’ (ABA), ‘response to auxin’, ‘response to salicylic acid’ (SA), ‘response to jasmonic acid’ (JA), ‘jasmonic acid mediated signalling pathway’, and ‘response to ethylene’ in the BPs and ‘indole-3-acetonitrile nitrilase activity’ in the molecular functions (MFs) were enriched (Fig. 6A). According to our GO analysis of the downregulated gene clusters, BP terms ‘water channel activity’, ‘glycerol channel activity’ and ‘nicotianamine synthase activity’ were enriched, and some cellular component (CC) terms about membrane and cell wall as well as ‘cellular water homeostasis’, ‘water transport’ and ‘response to water deprivation’ MF terms were enriched (Fig. 6B). The ‘biosynthesis of secondary metabolites’ KEGG pathway was significantly enriched in 169 upregulated DEGs, while no pathways were enriched in 59 downregulated DEGs (Fig. 6C). To study which genes responding to auxin, SA, JA and ABA were upregulated in the *CsFLC1* OE lines, we evaluated the transcript levels in the pathways mentioned above (Fig. 7). The results showed that almost all these genes had a higher expression level in the OE-*CsFLC1* lines than in the WT, but there were no significant differences in expression levels among the OE-*CsFLC1* lines. According to the transcriptome results, we noticed that three auxin-related genes, namely, *AUXIN RESPONSE FACTOR 5*/*MONOPTEROS* (*ARF5*/*MP*), *SENESCENCE-ASSOCIATED GENE 12* (*SAG12*) and the acyl acid amido synthetase gene *Gretchen Hagen 3*.*5* (*GH3*.*5*); three SA-related genes, namely, *THIONIN 2*.*1* (*THI2*.*1*), *MYB2* and *DIOXYGENASE 1* (*DOX1*); four JA-related genes, namely, *MYB57, TERPENE SYNTHASE* 03 (*TPS03*), *NAC055* and *VEGETATIVE STORAGE PROTEIN 1* (*VSP1*); and four ABA-related genes, namely, *DETOXIFICATION 48* (*DTX48*), *NAC92, CATALASE 1* (*CAT1*) and *BETA GLUCOSIDASE 18* (*BGLU18*), were significantly and obviously upregulated in the OE-*CsFLC1* lines.

**Fig. 6.**
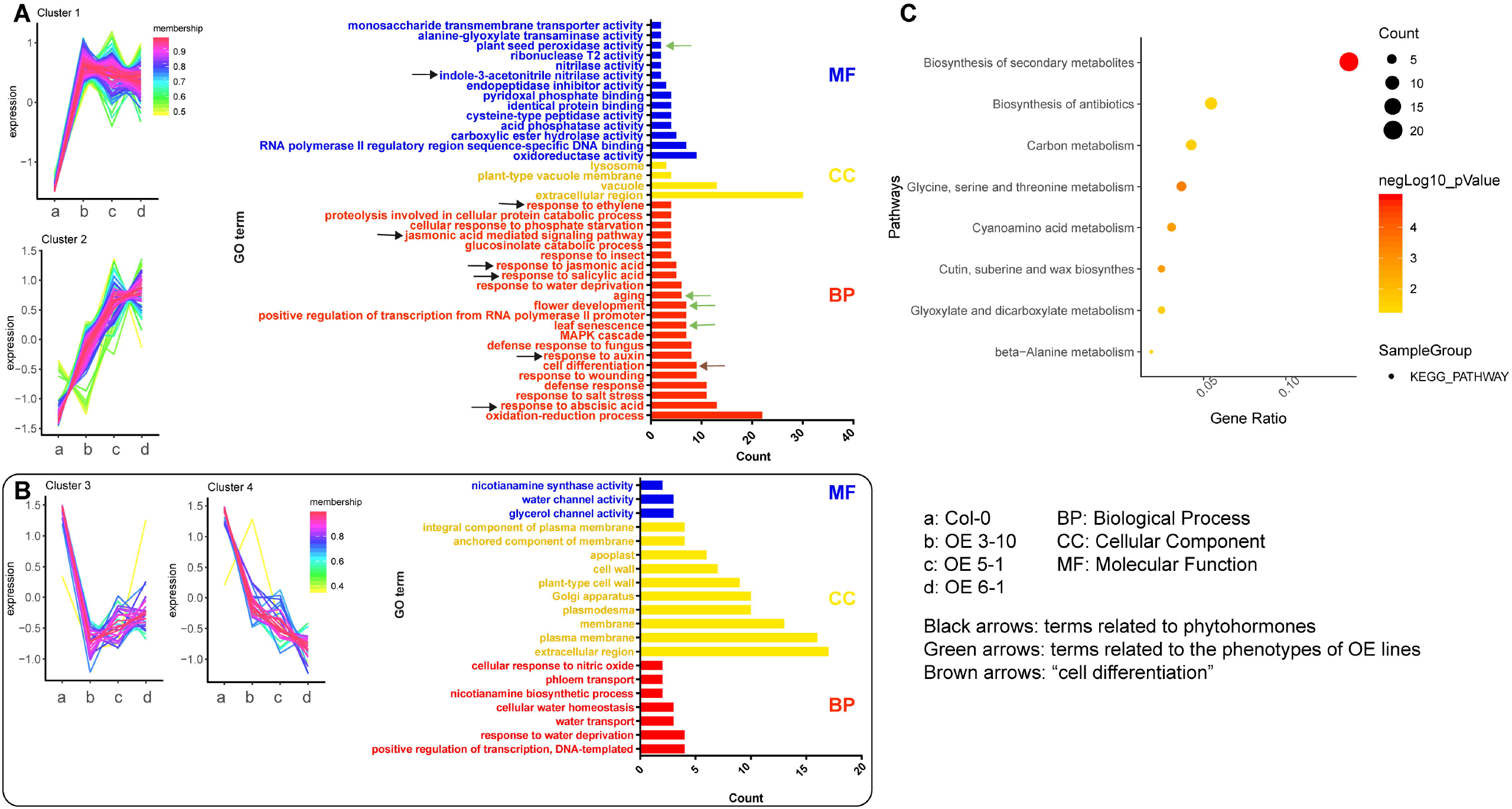
Transcriptome analysis of WT and *OE-CsFLC1* lines. (A) Gene cluster and GO enrichment of upregulated genes in *OE-CsFLC1* lines compared to WT. (B) Gene cluster and GO enrichment of downregulated genes in *OE-CsFLC1* lines compared to WT. (C) KEGG enrichment of upregulated genes in *OE-CsFLC1* lines compared to WT.

**Fig. 7.**
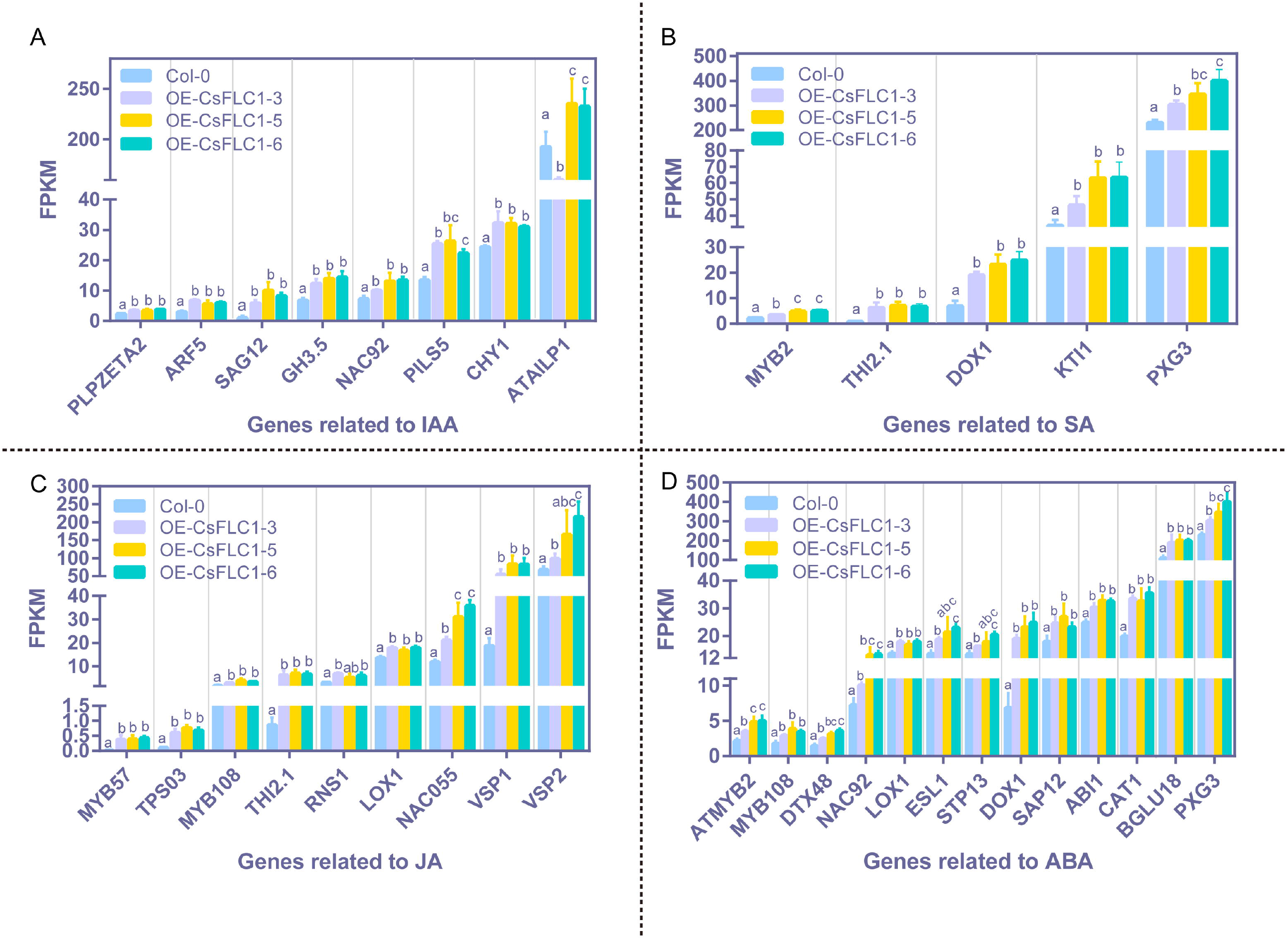
Expression of genes related to phytohormones in WT and *OE-CsFLC1* lines. (A) Expression of genes related to auxin in WT and *OE-CsFLC1* lines. (B) Expression of genes related to SA in WT and *OE-CsFLC1* lines. (C) Expression of genes related to JA in WT and *OE-CsFLC1* lines. (D) Expression of genes related to ABA in WT and *OE-CsFLC1* lines. SA: salicylic acid; JA: jasmonic acid; ABA: abscisic acid.

To explore which DEGs might be directly regulated by *CsFLC1* and cause the phenotypes of transgenic *Arabidopsis thaliana*, we analysed the expression patterns of known *AtFLC* direct target genes based on chromatin immunoprecipitation sequencing (ChIP-seq) data (Deng *et al*., 2011). In the OE *CsFLC1* lines, there were 13 common genes, among which 10 genes were induced, while 3 genes were repressed (Table S3). In *OE-CsFLC1* lines, four MADS-box family members, namely, *AGAMOUS-LIKE 42* (*AGL42*), *SEP3, SOC1* and *APETALA 3* (*AP3*); three hormone-related genes, namely, *BGLU18, NAC055* and *DTX48*; the posttranscriptional regulator *PUMILIO 8* (*APUM8*); the cytochrome P450 gene *CYP89A9*; and the wax biosynthesis-related gene *FATTY ACID REDUCTASE 3* (*FAR3*), were upregulated. However, *GUARD-CELL-ENRICHED GDSL LIPASE 18* (*GGL8*), the cell-wall-related gene *EXPA1* and the cytochrome P450 gene *CYP706A5* were downregulated (Fig. 8).

**Fig. 8.**
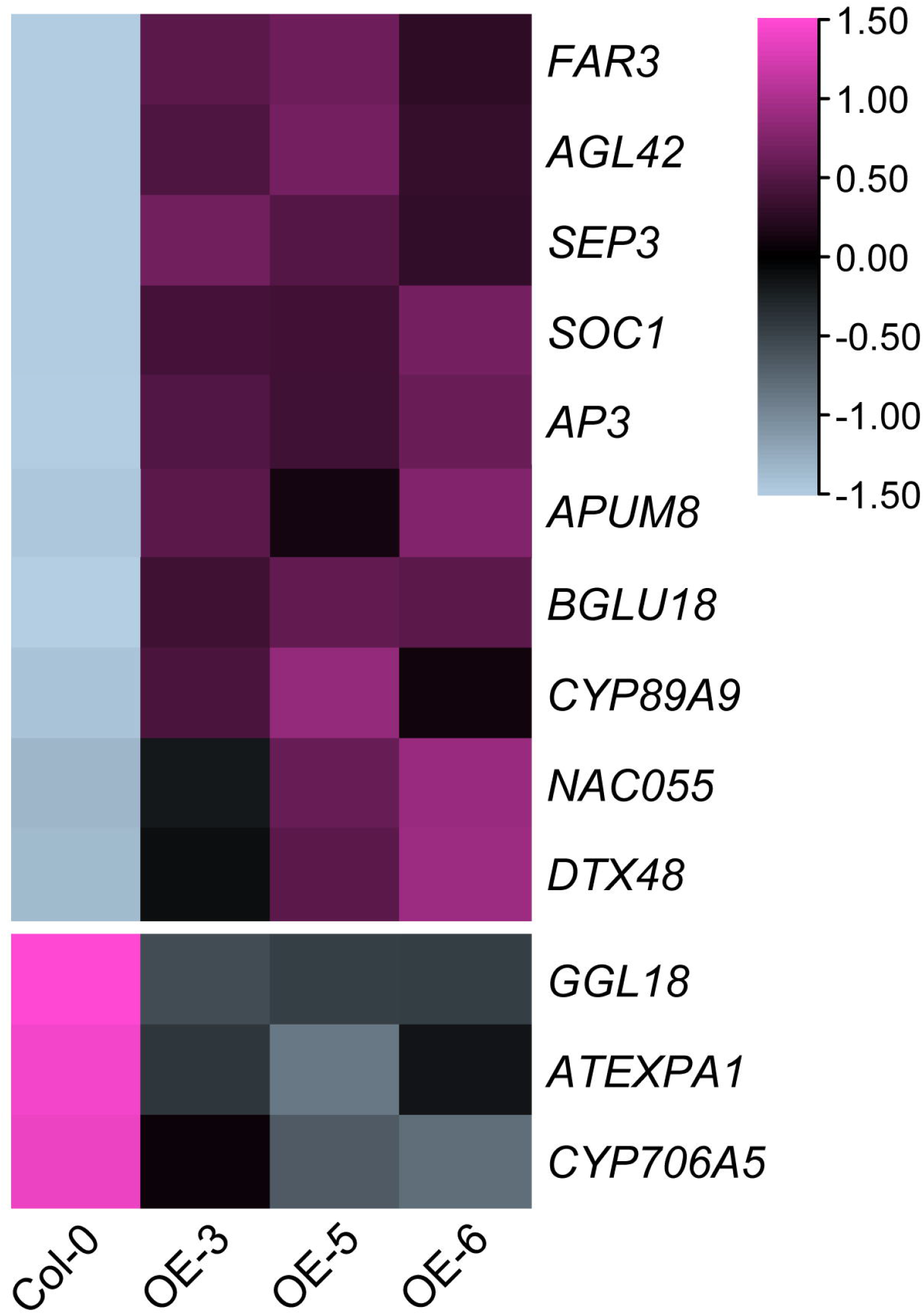
Expression patterns of *Arabidopsis* AtFLC target genes (Deng *et al*., 2011) in WT and *OE-CsFLC1* transgenic *Arabidopsis thaliana*. The pink colour represents upregulated genes, the blue colour represents downregulated genes, and the heatmap was generated by TBtools.

**Fig. 9.**
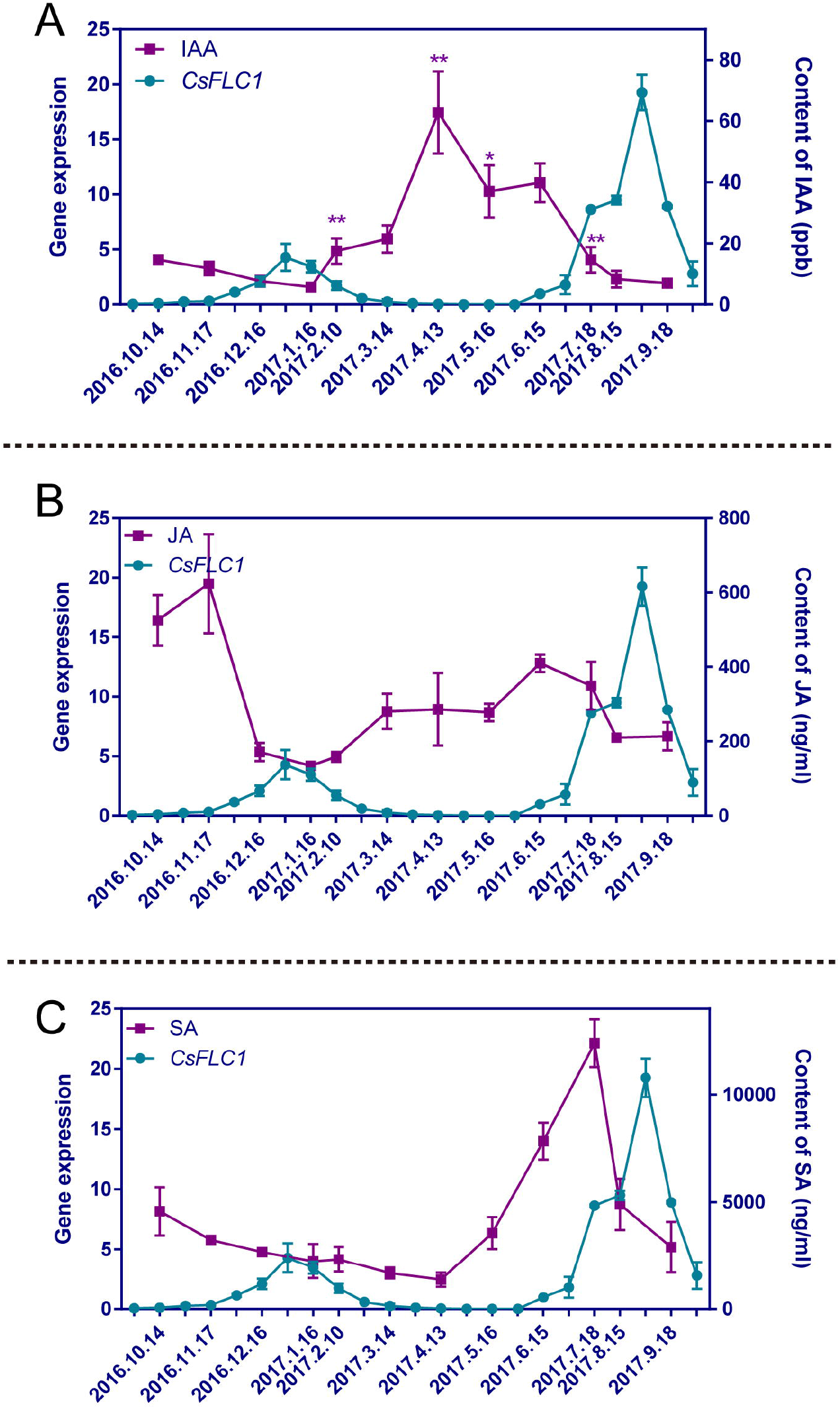
Variation trends of different phytohormone contents in tea plant during the whole year. (A) Content of IAA in tea plant throughout the year. (B) JA content of tea plant throughout the year. (C) Content of SA in tea plant throughout the year. IAA: indole-3-acetic acid; JA: jasmonic acid; SA: salicylic acid.

In conclusion, *CsFLC1* functions in controlling bolting and blooming times, seed germination, and leaf senescence as well as regulating some auxin-, SA-, JA- and ABA-associated genes in transgenic *Arabidopsis thaliana*.

### 3.5 Phytohormone contents in tea plant throughout the year

Since we found that there were some phytohormone pathways affected in the *35S::CsFLC1* lines, we measured the contents of three hormones (indole-acetic acid (IAA), JA and SA) in tea plant buds throughout the year. The hormone content changes are displayed in Fig. 7. The content of IAA was low from October 14^th^, 2016, to January 16^th^, 2017, and then increased until April 13^th^. The content peaked; later, it declined until July 18^th^, and finally, it was stable and low again (Fig 7A). The JA content was high on October 14^th^ and November 17^th^, decreased on December 16^th^, remained at a low level until February 10^th^, increased on March 14^th^, remained at a moderate level from February 10^th^ to May 16^th^, increased on June 15^th^, decreased until August 15^th^, and finally remained low until September 18^th^ (Fig. 7B). The content of SA was 4573.43 ng/ml on October 14^th^, 2016, slowly decreased until April 13^th^, 2017, rapidly increased until July 18^th^ and rapidly decreased until September 18^th^, sharply peaking on July 18^th^ (Fig. 7C). The changes in IAA and JA contents were opposite those of the *CsFLC1* expression level. There was only one time at which the SA content peak, which occurred one and-a-half months before the peak *CsFLC1* expression occurred, which was on August 28^th^. In conclusion, *CsFLC1* expression coincided with low contents of IAA and JA throughout the year but coincided with a high SA content of during flowering.

## 4. Discussion

### 4.1 Function of *CsFLC* in reproduction processes

Flowering time is regulated by five main pathways: the autonomous, gibberellin, ageing, photoperiod and vernalization pathways (Amasino and Michaels, 2010; He *et al*., 2020). These various pathways depend on different genetic regulations (Bloomer and Dean, 2017). *FLC* was identified first in 1999 and was considered to encode a repressor of flowering; its expression was upregulated by *FRIGIDA* (*FRI*) but downregulated by vernalization in Landsberg erecta (Ler; a late-flowering ecotype) (Michaels and Amasino, 1999; Sheldon *et al*., 1999). The *FLC* expression level was found to be an indicator of the extent of vernalization and was also regulated by the autonomous pathway (Sheldon *et al*., 2000; Michaels, 2001). *FLC* regulates circadian clock via autonomous and vernalization pathways to control flowering time, which shows that *FLC* is a link between vernalization and the circadian rhythm (Salathia *et al*., 2006). In addition, by binding to their chromatin, FLC represses the expression of *FT* and *SOC1* to inhibit flowering (Searle, 2006; Seo *et al*., 2009). In apple trees, the *MdFLC-like* gene is expressed in flower buds and is upregulated during cold accumulation and flower primordium differentiation and development (Soichiro *et al*., 2019).

In our previous study, we found that, during the flowering period, *CsFLC1* was specifically expressed during the floral transition stage (Liu *et al*., 2020). In this study, according to the expression patterns, *CsFLC1* was highly expressed during flower organ development. Thus, it positively regulated floral development, which coincided with the function of the apple *MdFLC-like* gene. Based on these findings, we constructed transgenic *CsFLC1* OE *Arabidopsis* in the Col-0 (an early-flowering ecotype) background. With respect to 35S::*CsFLC1* transgenic lines, bolting and blooming occurred earlier than it did for Col-0. According to the GO analysis of the transcriptome data, genes annotated to the ‘flower development’ term were enriched in the OE lines. We measured *CsFLC1* expression in the apical and axillary buds, flower buds and pistils of tea plant as well as in the carpel of *pCsFLC1::GUS::GUS* transgenic *Arabidopsis*. Our results showed that *CsFLC1* might positively regulate early flowering, floral transition, petal and pistil development reproduction processes. The transcriptome results showed that *SOC1* was upregulated in OE-*CsFLC1* lines. Thus, *CsFLC1* might induce flowering by influencing the expression of *SOC1* in tea plant, the results of which are opposite those from Ler *Arabidopsis* plants for both phenotype and genetic regulatory relationships. *AGL42*, a *SOC1*-*like* gene that was reported to be a target of *SOC1* and *FLC*, can promote flowering (Deng *et al*., 2011; Dorca-Fornell *et al*., 2011). *SEP3* is essential for floral meristem determinacy (Véronique *et al*., 2018). *AP3 is* involved in the formation of petals and stamens during flower development (Wuest *et al*., 2012). In our study, these four MADS-box genes *SOC1, AGL42, SEP3* and *AP3* were upregulated by *CsFLC1*, indicating that *CsFLC1* promotes flowering by controlling the expression of these genes directly in *Arabidopsis*.

### 4.2 *CsFLC1* putative function in bud dormancy

In tea plant, *CsFLC1* was highly expressed during the winter, which was identified as the endodormancy period of axillary buds (Hao *et al*., 2017). Based on the results of the GUS location patterns, *CsFLC1* was highly expressed in light under LDs, while it was induced in the dark under SDs, but it was highly expressed continuously under MDs. Interestingly, *CsFLC* accumulated in the day under LD conditions but accumulated at night under SD conditions (Fig. 3). This means that it plays different roles under different photoperiods. In previous studies, the transcript of *FLC* was independent of the photoperiod (Michaels and Amasino, 1999; Sheldon *et al*., 2000), indicating that *CsFLC1* is a special *FLC* with dual function in response to different photoperiods. Additionally, in the *CsFLC1::GUS::GUS* lines, *CsFLC1* could respond to low temperature. SDs and low temperature are the signals of winter, and plants enter dormancy when they receive the signals of winter to avoid damage and no longer grow before the environment is suitable for growth.

To further explore whether *CsFLC1* was coordinated with bud dormancy, we detected its expression in the late budbreak cultivar ZHDB from September 30^th^, 2016, to May 16^th^, 2017 (Fig. S1). The results showed that the mRNA transcript of *CsFLC1* were no longer detected approximately one month later than in LJ43, which coincided with budbreak time. Combining the results above, we predict that CsFLC1 is a repressor of bud break and maintains the bud dormancy state.

Relationships between *CsFLC1* expression and phytohormone concentrations and responses Phytohormones are important in the growth, development and response to environmental stimuli of plants. It has been reported that flowering and growth are promoted by relatively low concentrations of auxin but inhibited by higher concentrations (Leopold and Thimann, 1949). The results of whole-year RNA expression and hormone content detection showed that *CsFLC1* RNA transcription was correlated with a low content of IAA in both axillary and floral buds of tea plant. ARFs are TFs that bind to AuxREs in the promoters of early auxin response genes (Guilfoyle and Hagen, 2001; Tiwari *et al*., 2003). *ARF5* is an important activator in auxin signalling and is expressed in cells with low levels of auxin (Cucinotta *et al*., 2021). *SAG12* is a senescence indicator and is repressed by auxin (Noh and Amasino, 1999; Grbić, 2003; James *et al*., 2018). *GH3*.*5* functions in modulating and integrating both auxin and SA signalling (Staswick *et al*., 2002; Zhang *et al*., 2007), is expressed in seedlings, roots, stems, buds and blooming flowers of *Arabidopsis thaliana* (Zhang *et al*., 2008), and influences root development in rice (Fu *et al*., 2011). *ARF5, SAG12* and *GH3*.*5* were significantly upregulated in OE-*CsFLC1* lines, which indicated that *CsFLC1* might activate them to control auxin signalling and further induce flowering at low concentrations of auxin.

JA plays a critical role in inflorescence, stamen and seed development (Wasternack *et al*., 2013; Yuan and Zhang, 2015). JA induces the expression of *MYB57* to promote stamen filament development (Cheng *et al*., 2009). The JA content moderately peaked from May 16^th^ to August 15^th^ when the plants were in the floral transition and floral organ differentiation stages, and *MYB57* had an obviously higher transcription level in OE-*CsFLC1* lines. *NAC055* is a target of FLC and is induced by MeJA (Hickman *et al*., 2017); in our study, it was highly expressed in OE*-CsFLC1* transgenic *Arabidopsis*. These results showed that both *CsFLC1* and JA might control flowering through *NAC055*.

Cleland and Ajami (1974) found that SA could induce flowering in *Lemna gibba* G3. In this study, the content of SA increased during floral induction and initiation stages in tea plant and then decreased after July 18^th^. The results suggested that SA may play roles in floral induction of tea plant, which is in accordance with the results of previous studies on *Arabidopsis thaliana* (Martínez *et al*., 2004). *NAC055* could reduce the accumulation of SA (Zheng *et al*., 2012), and it was upregulated by *CsFLC1*. Therefore, the subsequent decrease in SA content might be because the high expression of *CsFLC1* induces the expression of *NAC055. DOX1* is expressed in the roots, anthers, and senescing leaves and induced by SA (De León *et al*., 2002). According to the results, the above *DOX1* expression level was increased in the OE-*CsFLC1* lines, which indicated that flowering induction and leaf senescence by *CsFLC1* and SA might occur through the *DOX1*-dependent signalling pathway.

ABA inhibits seed germination while accelerating floral transition and flower development (Giraudat, 1995; Hubbard *et al*., 2010; Fitzpatrick *et al*., 2011; Conti *et al*., 2014). ABA application can enhance *CAT1* expression in maize embryos (Guan *et al*., 2000). Based on the transcriptome results, *CAT1* expression was also significantly increased in the OE-*CsFLC1* lines. *MYB2* is a repressor of proanthocyanidins, and anthocyanin biosynthesis also plays a positive role in seed dormancy (Jun *et al*., 2015). Therefore, we proposed that OE-*CsFLC1* lines have defective seed germination, possibly because of the upregulation of ABA signalling genes, including *CAT1* and *MYB2*.

In conclusion, *CsFLC1* was found to be involved in flowering in *Arabidopsis thaliana* and tea plant, possibly by influencing flowering-related genes (*SOC1, AGL42, SEP3* and *AP3*) and hormone signalling, accumulation and metabolism.

## Abbreviations

ABA: Abscisic acid
AP3: APETALA3
ARF5/MP: AUXIN RESPOSE FACTOR 5/ MONOPTEROS
CAT1: CATALASE 1
CAL: CAULIFLOWER
CDS: Coding sequences
DEGs: Differentially expressed genes
DOX1: DIOXYGENASE 1
FT: FLOWERING LOCUS T
FLC: FLOWERING LOCUS C
GO: Gene ontology
GH3.5: Gretchen Hagen 3.5
IAA: Indoleacetic acid
JA: Jasmonic acid
KEGG: Kyoto Encyclopedia of Genes and Genomes
LD: Long day
MD: Middle day
MF: Molecular function
OE: Overexpression
SA: Salicylic acid
SAG12: SENESCENCE-ASSOCIATED GENE 12
SHP: SHATTERPROOF
SAMs: Shoot apical meristems
SD: Short day
SOC1: SUPPRESSOR OF OVEREXPRESSION OF CO 1
TPS03: TERPENE SYNTHASE 03
THI2.1: THIONIN 2.1
TF: Transcription factor
VSP1: VEGETATIVE STORAGE PROTEIN 1

## Supplementary data

Supplementary data are available at *JXB* online

Fig. S1 Expression pattern of *CsFLC1* in the late budbreak cultivar ZHDB.

Fig. S2 Counts of DEGs in *OE-CsFLC1* lines. A: Col-0, B: OE-3, C: OE-5, D: OE-6

Table S1. Accession numbers of the *SBP-, SEP3-* and *FLC-like* genes that form the clusters analysed in this study. In each cluster, the relative orientation of each gene is indicated by > or <.

Table. S2 Sequences of primer pairs used in this paper.

Table S3 Expression patterns of *AtFLC* target genes in WT and *OE-CsFLC1* transgenic *Arabidopsis thaliana*.

## Acknowledgements

This work was supported by the National Key Research and Development Program of China (2018YFD1000601), the Major Science and Technology Special Project of Variety Breeding of Zhejiang Province (2016C02067), the Agricultural Science and Technology Innovation Program of the Chinese Academy of Agricultural Sciences (CAAS-ASTIP-2021-TRICAAS) and the China Agriculture Research System of MOF and MARA (CARS-19). We would like to thank Professor Cristina Ferrándiz from the Spanish National Research Council for advice on revising the manuscript.

## Author contributions

YL, XH and XW conceived, designed and supervised the experiment. YL, HZ, XZ, NL, KZ, TD, LW and YY conducted the experiments, which included collecting the samples, measuring the phytohormone contents and extracting RNA. YL and XW analysed the data. LD contributed to the evolutionary analysis. YL, LD and XW wrote the manuscript.

## Data availability statement

## Figure legends

**Fig. S1 Expression pattern of *CsFLC1* in the late budbreak cultivar ZHDB**.

**Fig. S2 Counts of DEGs in *OE-CsFLC1* lines**. A: Col-0, B: OE-3, C: OE-5, D: OE-6.

**Table S1. Accession numbers of the *SBP-, SEP3-* and *FLC-like* genes that form the clusters analysed in this study**. In each cluster, the relative orientation of each gene is indicated by > or <.

**Table. S2 Sequences of primer pairs used in this paper**.

**Table S3 Expression patterns of *AtFLC* target genes in WT and *OE-CsFLC1* transgenic *Arabidopsis thaliana***.

